# Dynein-2 intermediate chains play crucial but distinct roles in primary cilia formation and function

**DOI:** 10.1101/251694

**Authors:** Laura Vuolo, Nicola L. Stevenson, Kate J. Heesom, David J. Stephens

## Abstract

The dynein-2 microtubule motor is the retrograde motor for intraflagellar transport. Mutations in dynein-2 components cause skeletal ciliopathies, notably Jeune syndrome. Dynein-2 comprises a heterodimer of two non-identical intermediate chains, WDR34 and WDR60. Here, we use knockout cell lines to demonstrate that each intermediate chain has a distinct role in cilia function. Both proteins are required to maintain a functional transition zone and for efficient bidirectional intraflagellar transport, only WDR34 is essential for axoneme extension. In contrast, only WDR60 is essential for co-assembly of the other subunits. Furthermore, WDR60 cannot compensate for loss of WDR34 or vice versa. This work defines a functional asymmetry to match the subunit asymmetry within the dynein-2 motor. Analysis of causative point mutations in WDR34 and WDR60 can partially restore function to knockout cells. Our data show that Jeune syndrome is caused by defects in transition zone architecture as well as intraflagellar transport.

**SUMMARY:** Here, Vuolo and colleagues use engineered knockout human cell lines to define roles for dynein-2 intermediate chains. WDR34 is required for axoneme extension, while WDR60 is not. Both subunits are required for cilia transition zone organization and bidirectional intraflagellar transport.

## Introduction

Cytoplasmic dyneins are minus-end directed motors that use the energy of ATP hydrolysis to move along microtubules. Two cytoplasmic dyneins have been identified. The better-characterized dynein-1 is involved in the transport of cargos in the cytoplasm, organelle dynamics and in mitotic spindle organization during mitosis (Roberts et al., 2013). Dynein-2 is responsible for retrograde transport in cilia and flagella. Primary (non-motile) cilia are hair-like extensions present on almost all animal cells that act as antennae for extracellular signals and are fundamental to proper metazoan development and ongoing health. They integrate signals in key pathways including sonic hedgehog (Shh), Wnt and platelet-derived growth factor signaling and participate in metabolic control and autophagy (Reiter and Leroux, 2017). Cilia are particularly important to ensure correct Shh signaling during embryonic development (Goetz and Anderson, 2010; He et al., 2017). Defects in cilia are linked to many human diseases, known collectively as ciliopathies, including developmental disorders, neurodegeneration and metabolic diseases (Reiter and Leroux, 2017; Yee and Reiter, 2015).

Ciliogenesis is initiated in non-dividing cells by the docking of pre-ciliary vesicles with the mother centriole. The pre-ciliary vesicles fuse and then surround the mother centriole concomitant with the assembly of a series of protein modules that form a diffusion barrier separating the distal end of the mother centriole from the rest of the cell cytoplasm (Garcia-Gonzalo et al., 2011). A microtubule bundle, the axoneme, then extends from the centriole to allow cargo transport by the process of intraflagellar transport (IFT) (Yee and Reiter, 2015). A “transition zone” (TZ) that separates the mother centriole from the main length of the axoneme forms a diffusion barrier for both soluble and membrane proteins at the base of the cilium (Garcia-Gonzalo et al., 2011; Garcia-Gonzalo and Reiter, 2017). Once established, cilia are maintained and operated by the process of IFT (Hou and Witman, 2015). IFT-B complexes (comprising of a core subcomplex, including nine subunits (IFT88, -81, -74, -70, -52, -46, -27, -25, and -22) with five additional, peripherally-associated subunits (IFT172, -80, -57, -54, and -20)) undergo kinesin-2-driven motility from base to tip where the complexes are then reorganized prior to retrograde transport of IFT-A (comprising 6 subunits (IFT144, IFT140, IFT139, IFT122, IFT121/WDR35, and IFT43)) driven by dynein-2 (Hou and Witman, 2015; Jensen and Leroux, 2017). This process ensures the correct localization of receptors and signaling molecules within cilia as well as act in signal transduction from the cilium to the rest of the cell.

The genes encoding subunits of the dynein-1 and dynein-2 motors are largely distinct. Some light chain subunits are common to both but the major subunits (heavy, intermediate and light intermediate chains) are different between the two holoenzyme complexes. Dynein-2 is built around a heavy chain dimer of DYNC2H1 (DHC2) (Criswell et al., 1996; Mikami et al., 2002). This associates with two intermediate chains, WDR34 and WDR60, first identified as dynein-2 subunits named FAP133 and FAP163 in *Chlamydomonas* (Patel-King et al., 2013; Rompolas et al., 2007) and subsequently shown to be components of metazoan dynein-2 (Asante et al., 2013; Asante et al., 2014). This asymmetry distinguishes dynein-2 from dynein-1 where two identical IC subunits form the holoenzyme. The reason for this asymmetry is unclear. In addition, a dynein-2-specific light intermediate chain (DYNC2LI1/LIC3) has been identified (Hou and Witman, 2015; Mikami et al., 2002) as well as a specific light chain, TCTEX1D2 (Asante et al., 2014; Schmidts et al., 2015). Mutations in genes encoding dynein-2 subunits are associated with skeletal ciliopathies, notably short rib-polydactyly syndromes (SRPSs) and Jeune asphyxiating thoracic dystrophy (JATD, Jeune syndrome). These are recessively inherited developmental disorders characterized by short ribs, shortened tubular bones, polydactyly and multisystem organ defect (Huber and Cormier-Daire, 2012). In recent years, whole exome-sequencing technology has enabled the identification of new mutations involved in skeletal ciliopathies, notably a range of mutations affecting dynein-2 heavy chain (DYNC2H1 or DHC2, (Chen et al., 2016; Cossu et al., 2016; Dagoneau et al., 2009; El Hokayem et al., 2012; Mei et al., 2015; Merrill et al., 2009; Okamoto et al., 2015; Schmidts et al., 2013a)). Additionally, mutations in WDR34 (Huber et al., 2013; Schmidts et al., 2013b), WDR60 (Cossu et al., 2016; McInerney-Leo et al., 2013), LIC3 (Kessler et al., 2015; Taylor et al., 2015) and TCTEX1D2 (Schmidts et al., 2015) have also been reported. The role of dynein-2 heavy chain has been extensively studied in *Chlamydomonas, C. elegans*, and mice. In all cases, loss of dynein heavy chain results, in short, stumpy cilia that accumulate IFT particles at the tip, consistent with a role of dynein-2 in retrograde ciliary transport (Hou and Witman, 2015). Recently, more interest has been focused on the role of the subunits associated with DHC2. Two studies in *Chlamydomonas* and in human patient-derived fibroblasts revealed that LIC3 (D1bLIC in *Chlamydomonas)* plays an important role for ciliogenesis and stability of the entire dynein-2 complex (Li et al., 2015; Taylor et al., 2015). Similarly, loss of Tctex2b (TCTEX1D2) destabilizes dynein-2 and reduces IFT in *Chlamydomonas* (Schmidts et al., 2015).

Previous work from our lab and others has shown that loss of function of dynein-2 intermediate chains, WDR34 and WDR60, is associated with defects in cilia. Knockdown of WDR60 or WDR34 in hTERT-RPE1 cells results in a reduction of ciliated cells, with an increase or decrease of the cilia length, likely depending on depletion efficiency (Asante et al., 2014). Mutations in WDR34 have also been shown to result in short cilia with a bulbous ciliary tip in patients fibroblast cells affected by SRP (Huber et al., 2013). Consistent with the results obtained in patient cells, loss of WDR34 in mice also results in short and stumpy cilia with an abnormal accumulation of ciliary proteins and defects in Shh signaling (Wu et al., 2017). Similarly, mutations in WDR60 patient fibroblasts are associated with a reduction in cilia number, although the percentage of ciliated cells was variable in different affected individuals (McInerney-Leo et al., 2013). These findings are all consistent with roles for WDR34 and WDR60 in IFT.

In this study, we sought to better understand the role of dynein-2 in human cells using engineered knockout (KO) cell lines for WDR34 and WDR60. We define a functional asymmetry within the complex, where WDR34 is absolutely required for ciliogenesis, while WDR60 is not. In contrast, WDR60 is essential to maintain the integrity of the ciliary transition zone and for retrograde trafficking of IFT particles. Furthermore, by expressing HA-tagged WDR34 in WDR60 KO cells and HA-tagged WDR60 in WDR34 KO cells, we found that WDR34 is not required for the other subunits to assemble, whereas loss of WDR60 leads to significant defects in dynein-2 holocomplex assembly. We propose a model where dynein-2 requires WDR34 for axoneme extension but not for the assembly of the other subunits of the complex, whereas WDR60 is crucial for dynein-2 stability, IFT, and ciliary transition zone assembly and/or maintenance. Analysis of disease-causing patient mutations further defines the role of dynein-2 in cilia formation and function.

## Results

### WDR34 or WDR60 play different roles in cilia function

To understand the function of WDR34 and WDR60, we generated KO human telomerase-immortalized RPE1 (hTERT-RPE1) cells using CRISPR-Cas9. We derived two WDR34 KO clones (1 and 2) using guide RNAs (gRNAs) targeting exons 2 and 3, and one KO clone for WDR60, targeting exon 3. Genomic sequencing of these clones identified insertion/deletion mutations on the targeted sequences (Fig. S1). All cell clones were analyzed for protein expression by immunoblot using polyclonal antibodies. Neither WDR34 nor WDR60 was detected in the respective KO cells compared to the controls (Fig. S2A and Fig. S2B) which provides evidence that downstream initiation sites are not being used. To mitigate against the possibility of any off-target effects, we grew KO cells alongside control CRISPR cells which had been transfected and treated the same way as the KO, but genomic sequencing showed no mutation at the target site. These cells (WDR34 KO CTRL and WDR60 KO CTRL) did not present any cilia defects when stained with Arl13b or IFT88 (Fig. S2C and S2D). Images in all figures show WT cells where indicated. Indistinguishable results were obtained using these control cell lines. Defects in ciliogenesis in both WDR34 and WDR60 KO cells were rescued by overexpressing WT proteins, confirming that the phenotypes we observed were not due to off-target mutations (described below).

Loss of WDR34 severely impaired the ability of these cells to extend a microtubule axoneme (Fig. 1A, B), although Arl13b localized within those few cilia that did form. In contrast, loss of WDR60 did not significantly affect the ability of cells to extend an axoneme (Fig. 1B). Cilia were shorter in both WDR60 and WDR34 KO cells (Fig. 1C). Next, we examined the assembly and structure of primary cilia in WDR34 and WDR60 KO cells by transmission electron microscopy (EM). After 24 hr of serum starvation, WT RPE1 cells extend a defined axoneme surrounded by a ciliary membrane (Fig. 1D). In contrast, WDR34 KO cells failed to extend an axoneme (Fig. 1E) but showed a large docked pre-ciliary vesicle, consistent with the small Arl13b-positive structures seen by immunofluorescence. WDR60 KO cells showed apparently normal cilia (Fig. 1F) with normal basal body structures and axoneme extension. However, when an entire cilium was captured in WDR60 KO serial sections (Fig. 1G), we observed a bulged cilia tip containing accumulated electron dense particles (Fig. 1H). To our surprise, we also observed the ciliary membrane bulged at a second point along the axoneme and this region contained intraciliary vesicular structures (Fig. 1Hi).

**Figure 1:**
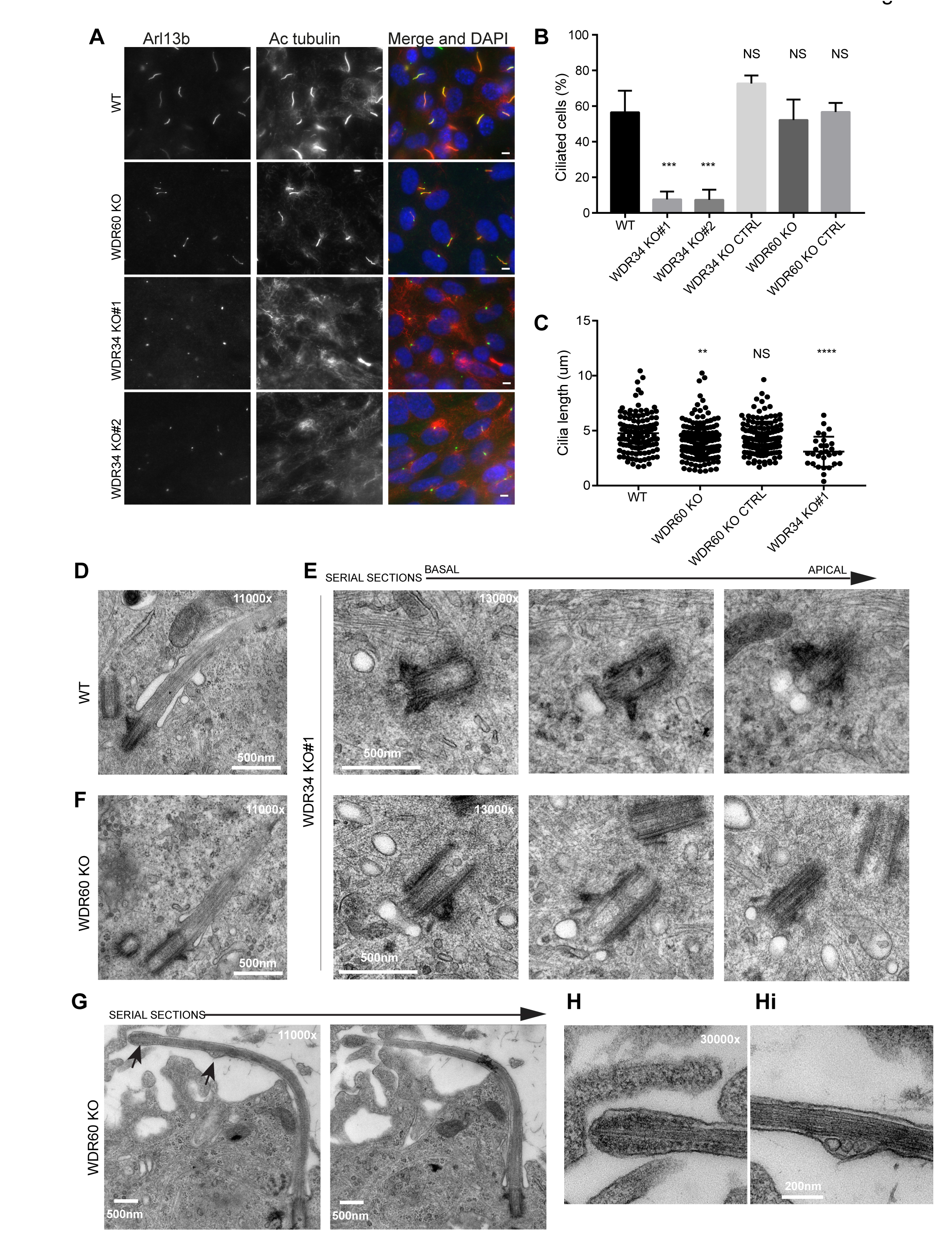
Role of dynein-2 intermediate chains WDR34 and WDR60 in ciliogenesis. (A) Cilia stained with the markers Arl13b (green) and acetylated tubulin (AcTub, red) in RPE1 WDR60 and WDR34 KO cell lines. Scale bars 5µ.m. (B) Percentage of ciliated cells (n=3; 656 WT, 430 WDR34 KO#1, 296 WDR34 KO#2, 397 WDR34 KO CTRL, 343 WDR60 KO and 195 WDR60 KO CTRL cells quantified). (C) Cilia length in WDR60 and WDR34 KO compared with WT cells and CRISPR control cells lines (n=3; 120 WT, 158 WDR60 KO, 138 WDR60 KO CTRL and 30 WDR34#1 cells quantified). Mann-Whitney test was used, p-value: ****=<0.0001. (D-Hi) Representative 70 nm thick EM sections of (D) WT, (E) WDR34 KO and (F, G-Hi) WDR60 KO cilia. (E) Six serial sections through a WDR34 KO cilium showing no axoneme extension. (G) Two serial sections through a WDR60 KO cilium showing. Arrows point to the bulbous ciliary tip and to a membrane protrusion containing membrane vesicles; enlargements are shown to the right (H and Hi). Scale bar length and magnification is indicated on the images.

### Loss of WDR34 and WDR60 causes accumulation of proteins at the ciliary tip

The abnormal structure of cilia in the KO cells led us to analyze the steady-state localization of the IFT machinery. After 24 hr serum starvation, IFT88 (part of IFT-B) was found almost exclusively at the base of the cilia in wild-type (WT) RPE1 cells (Fig. 2A, quantified in Fig. 2Ai), but in WDR60 KO and WDR34 KO cells IFT88 was found throughout the cilia and accumulated at the tip (Fig. 2A, 2Aii, and 2Aiii). Similarly, another IFT-B component, IFT57 (Fig. 2B) was enriched at the cilia tips in WDR60 and WDR34 KO cells. Quantification of the localization of these IFT-B proteins showed an accumulation of both IFT88 (Fig. 2C) and IFT57 (Fig. 2D) at both the base and tip of the cilia in WDR60 KO cells. In WDR34 KO cells, IFT-B proteins were seen to accumulate at the tip of cilia. The accumulation at the base seen in WDR60 KO cells was not evident in WDR34 KO cells. The limited numbers of ciliated WDR34 KO cells precluded further quantification. Another IFT-B protein, IFT20 (Fig. 2E), was enriched at both the tip and the base of cilia in WDR60 KO cells (quantification in Fig. 2F). IFT20 is the only IFT component found to localize to the Golgi until now. As expected, IFT20-GFP was found at the Golgi in WT cells, although the Golgi pool of IFT20-GFP seen in WT cells was largely absent from WDR60 KO cells (Fig. 2G).

**Figure 2:**
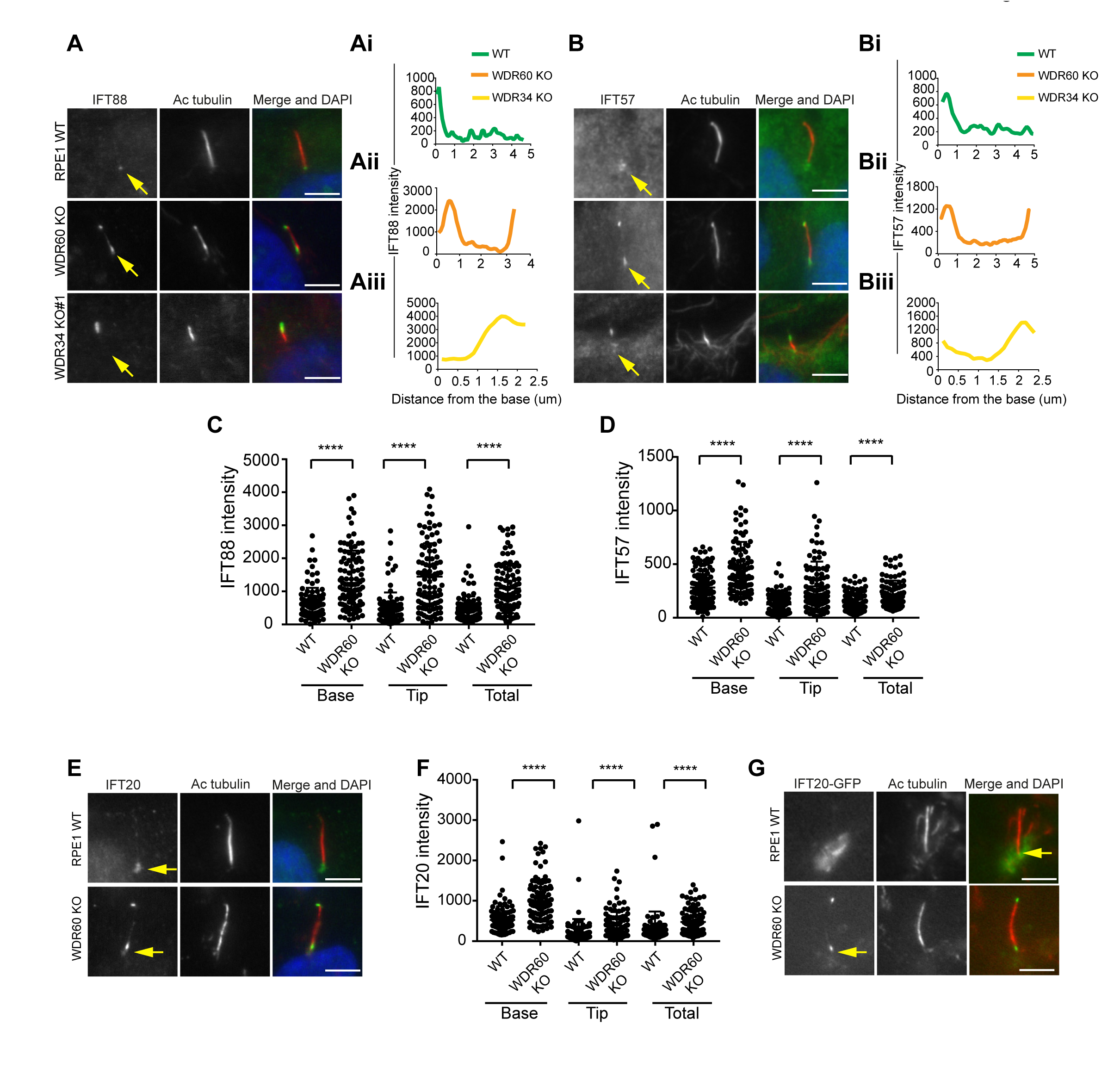
WDR34 and WDR60 are essential for IFT-B trafficking in primary cilia. (A, B) Localization of (A) IFT88 and (B) IFT57 in WT, WDR60 KO, and WDR34 KO#1 cells. Line graphs show lines scans of IFT intensity along the length of a representative cilium from WT (Ai and Bi, green), WDR60 KO (Aii and Bii, orange) and WDR34 KO (Aiii and Biii, yellow) cells. (C, D) Quantification of IFT-B localization within cilia in WT and WDR60 KO cells (C, IFT88 102 WT and 101 WDR60 cells quantified; D, IFT57 106 WT and 98 WDR60 KO cells quantified; n=3 independent experiments). Mann-Whitney test was used, p-value: ****=<0.0001. (E) Endogenous IFT20 accumulates at the ciliary tip in WDR60 KO cells. (F) Quantification of IFT20 localization within cilia in WDR60 KO cells (n=3 independent experiments, 102 WT and 100 WDR60 KO cells quantified). (G) Localization of IFT20-GFP in WT and WDR60 KO fixed cells. Scale bar, all panels = 5 µm. Arrows point to the ciliary base.

Next, we analyzed the localization of IFT-A proteins. We found that both IFT140 (Fig. 3A) and IFT43 (Fig. 3B) were accumulated along the length of the axoneme within cilia in WDR60 KO cells, as well as at the tips in the few cilia present in WDR34 KO cells, while they were found only at the base of the cilia in WT cells (quantification in Fig. 3Ai-iii and 3Bi-iii). Further quantification showed that both IFT140 (Fig. 3C) and IFT43 (Fig. 3D) are accumulated in WDR60 KO cilia compared to controls. Both IFT140 and IFT43 show a more pronounced accumulation at ciliary tips in WDR34 KO cells. In addition, we determined the localization of a subunit of the anterograde kinesin-2 motor, KAP3 which was also accumulated at the ciliary tip of WDR60 KO (Fig. 3E).

**Figure 3:**
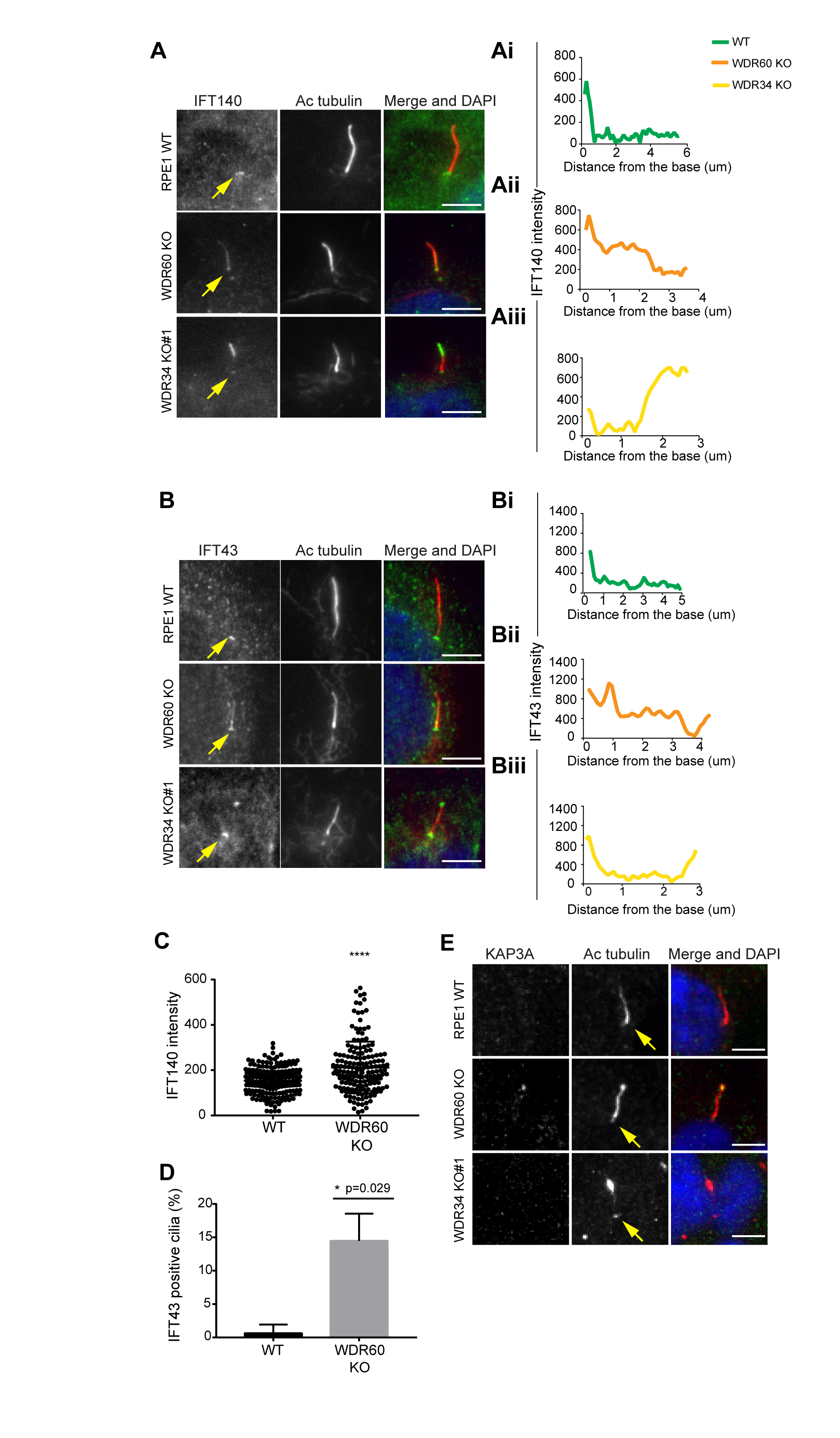
IFT-A trafficking defects in absence of WDR34 and WDR60. (A-B) Localization of IFT-A proteins (A) IFT140 and (B) IFT43 in WT, WDR60 KO, and WDR34 KO#1 cells. Line graphs show lines scans of IFT intensity along the length of a representative cilium from WT (Ai and Bi, green), WDR60 KO (Aii and Bii, orange) and WDR34 KO (Aiii and Biii, yellow) cells. (C) Quantification of IFT140 intensity within cilia in WT and WDR60 KO cells (n=3, 186 WT and 166 WDR60 KO cells quantified). (D) Quantification of the number of IFT43 positive cilia from WT and WDR60 KO cells (n=3, 271 WT and 203 WDR60 KO cells quantified). (C-D) Mann-Whitney test was used, p-value: ****=<0.0001. (E) Localization of KAP3A in WT, WDR60 KO, and WDR34 KO cells. Scale bars = 5µm. Arrows point to the ciliary base.

To study whether defects in dynein-2 affect the transport of membrane proteins we used GFP-fusions with Arl13b, somatostatin receptor type 3 (SSTR3), 5-hydroxytryptamine receptor type 6 (5HT6) and Rab8a. We found that in live cells Arl13b-GFP and EGFP-SSTR3 accumulate at the ciliary tip (Fig. 4A and 4B). Surprisingly, we also noticed a consistent reduction in the amount of Arl13b-GFP within cilia in WDR60 KO cells (Fig. 4C and 4Ci). The same observation was made with EGFP-SSTR3 (Fig. 4D and 4Di) and EGFP-5HT6 (Fig. 4E and 4Ei). In contrast, GFP-Rab8a localization in the cilia was indistinguishable between WT and WDR60 KO cells (Fig 4F and 4Fi).

**Figure 4:**
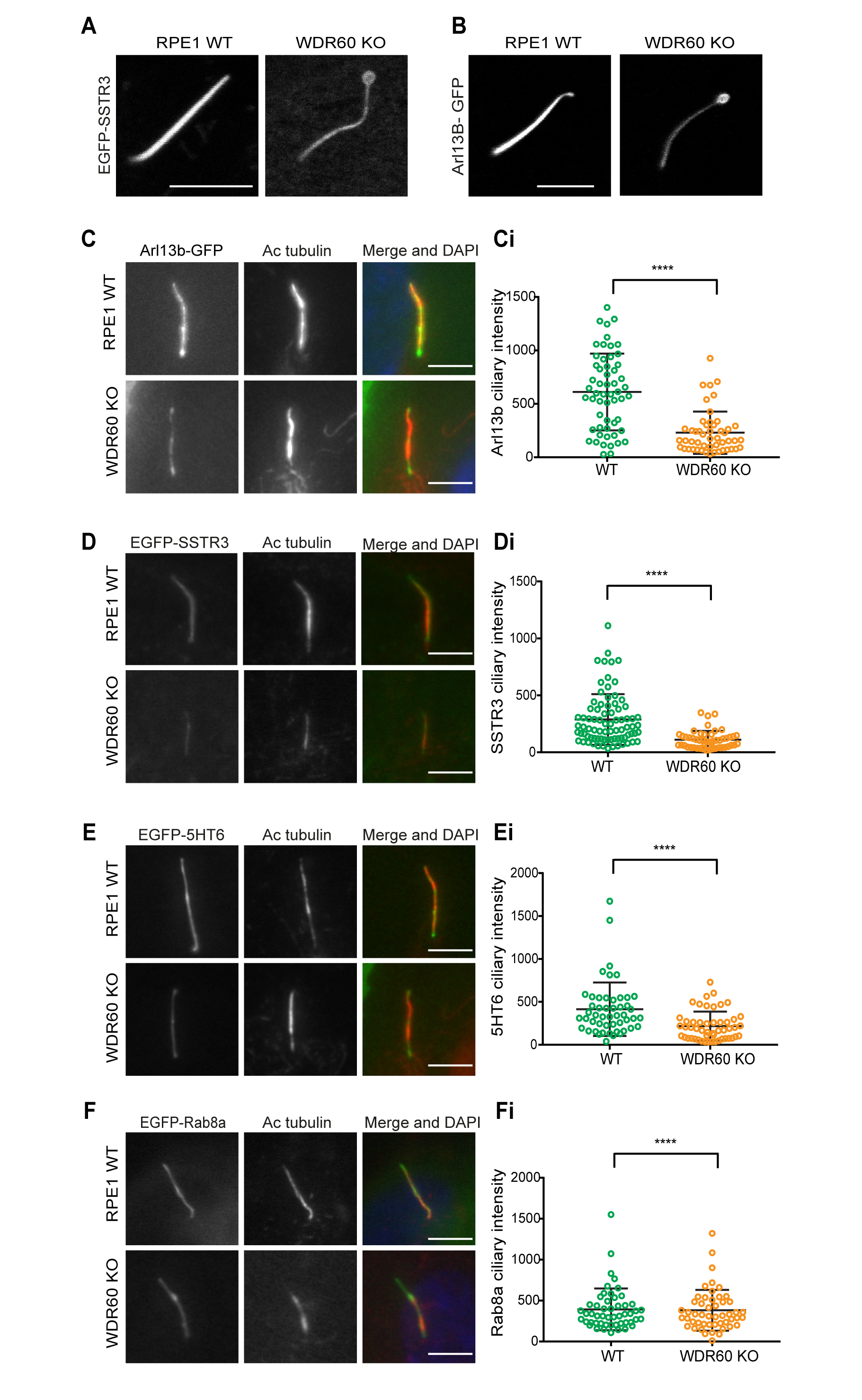
WDR60 is crucial for the composition of cilia membrane proteins. (A and B) Single frame images are taken from live imaging movies of WT and WDR60 KO cells overexpressing EGFP-SSTR3 and GFP-Arl13b. (C-F) Fixed cell staining of overexpressed Arl13b-GFP, EGFP-SSTR3, EGFP-5HT6, and EGFP-Rab8a in WT and WDR60 KO cells. (Ci-Fi) Intensity quantification of the overexpressed protein indicated (Arl13b-GFP n=3, 56 WT and 50 WDR60 KO cells quantified; EGFP-SSTR3 n=3, 80 WT and 50 WDR60 KO cells quantified; EGFP-5HT6 n=3, 50 WT and 51 WDR60 KO cells quantified; EGFP-Rab8a n=3, 51 WT, and WDR60 50 KO cells quantified). Mann-Whitney test was used, p-value: ****=<0.0001. Scale bars 5µ.m.

### Dynein-2 is required for ciliary transition zone assembly

Previous studies have shown that the protein content of cilia is maintained by a diffusion barrier formed by the transition zone. Changes in transition zone composition have been associated with the mislocalization of ciliary proteins, including the membrane marker Arl13b (Li et al., 2015; Shi et al., 2017). To test if the reduction in Arl13b seen in our WDR60 KO was caused by a defect in the transition zone, we labeled KO and WT cilia with known transition zone markers. We found that the core transition zone marker, RPGRIP1L (also known as MKS5), is no longer restricted to an area adjacent to the mother centriole in WDR60 KO cells (Fig. 5A, quantified in 5Ai). Conversely, TMEM67 (also known as MKS3), which in WT cilia extends from the basal body through a more distal region, becomes much more tightly restricted to the base of the cilium in WDR60 KO cells (Fig. 5B, quantified in 5Bi). We also determined the transition zone organization in WDR34 KO cells. The few cilia found in the WDR34 KO cells recapitulate the same phenotype observed in the WDR60 KO cilia with an expansion of RPGRIP1L to a more distal position and a reduction of the TMEM67 domain (Fig. 5A and Fig. 5B). In contrast to TMEM67 and RPGRIP1L, no changes were observed for the transition zone marker TCTN1 in both WDR34 KO and WDR60 KO cells with respect to the control (Fig. 5C).

**Figure 5:**
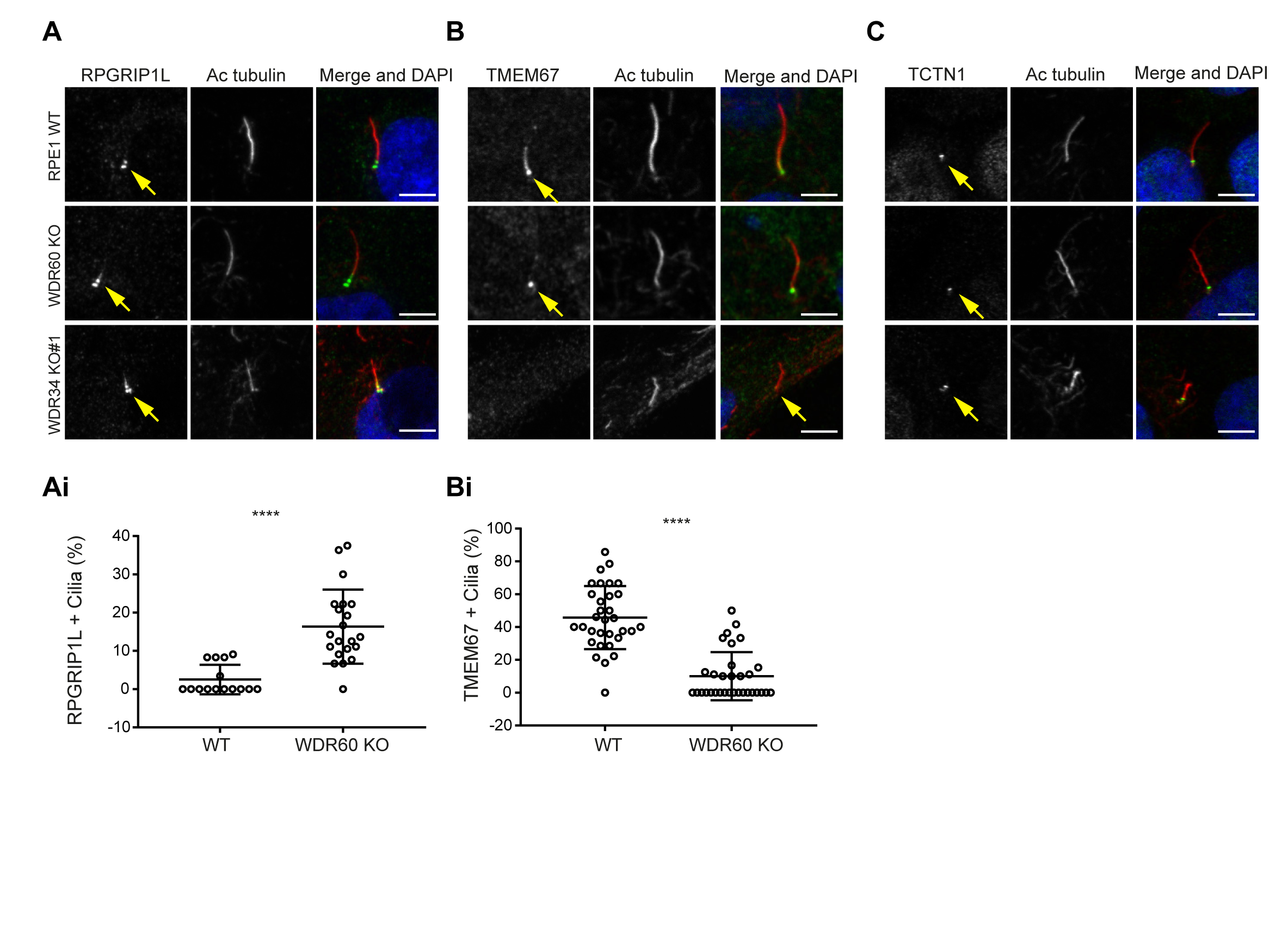
Dynein-2 is important for transition zone assembly. (A-C) Localization of RPGRIP1L, TMEM67, and TCTN1 in WT and KO cells. (Ai and Bi) Percentage of RPGRIP1L and TMEM67 positive cilia. RPGRIP1L n=3, 188 WT and 272 WDR60 KO cells quantified; TMEM67 n=3, 359 WT and 243 WDR60 KO cells quantified. Mann-Whitney test was used p-value: ****=<0.0001. Scale bars 5µ.m. Arrows point to the ciliary base.

### Unregulated entry of Smoothened into cilia following loss of WDR60

Defects in the dynein-2 motor have been previously associated with deregulation of the Shh pathway (May et al., 2005). Smoothened (Smo), a key component of Shh signaling, localizes to the cilia in response to Shh stimulation but is normally excluded from cilia in cells that have not been treated with Shh or an equivalent agonist such as Smoothened agonist (SAG). Unexpectedly, we found that Smo was localized to cilia in WDR60 KO cells even in the absence of SAG stimulation (Fig. 6A, and 6Ai). This localization did not increase upon agonist treatment. In contrast, Smo was indeed excluded from cilia in WT cells at steady state (Fig. 6A and 6Ai) but was readily detected within cilia following SAG stimulation (Fig. 6B, 6Bi, and 6Bii).

**Figure 6:**
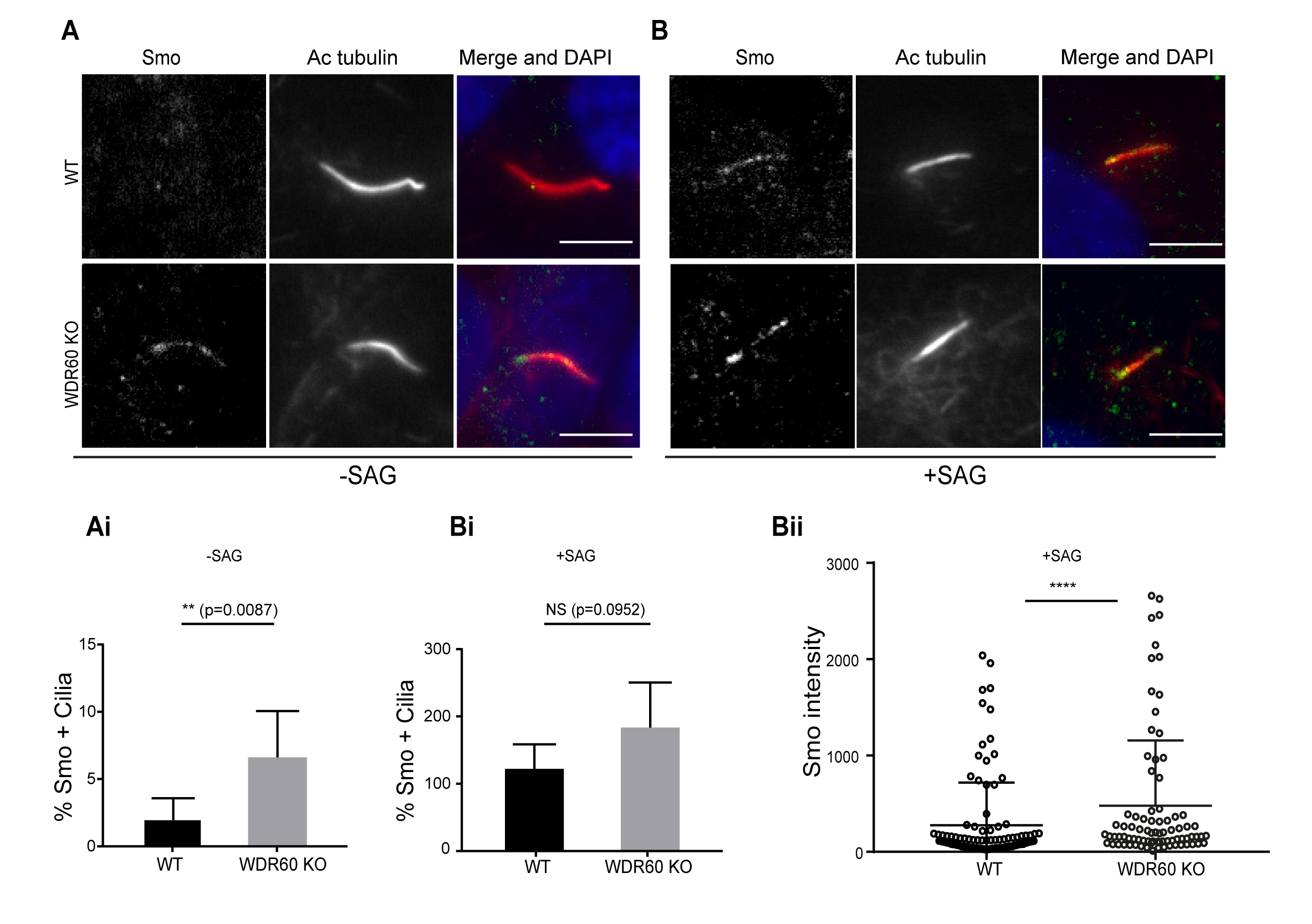
Loss of WDR60 affects Smo localization in the cilia. (A and B) Immunofluorescence of WT and WDR60 KO cells in presence or absence of SAG and stained for Smo (green), AcTub (red), and DAPI (blue). (Ai and Bi) Percentage of Smo positive cilia in SAG untreated (n=3, 148 WT and 120 WDR60 KO cells quantified) and treated cells (n=3, 670 WT and 580 WDR60 KO cells quantified). (Bii) Quantification of the total intensity of ciliary Smo labeling in cells treated with SAG (n=3, 102 WT and 82 WDR60 KO cells quantified). Mann-Whitney test was used, p-value: ****=<0.0001. Scale bars 5µ.m.

### Expression of wild-type and patient mutants of WDR60 and WDR34 in KO cells

Many mutations in WDR34 and WDR60 have been associated with SRPs and JATD syndromes. We engineered selected patient mutations into WDR34 and WDR60 to determine the defect compared to expression of wild-type proteins. Two WDR60 mutations were selected from one SRPS patient with compound heterozygosity for WDR60 (WDR60[T749M], WDR60[Q631*]) and two WDR34 mutations from two patients with JATD syndromes (WDR34[A22V], WDR34[C148F]). The first WDR60 mutation is located in the third WD repeat (WDR60[T749M]) and the second (WDR60[Q631*]) is located just before the WD repeat domain (Fig. S3A). We undertook “rescue” experiments in which we stably expressed WT or mutant versions of WDR34 and WDR60 in KO cells. Protein expression of stably transfected WT and HA-WDR60 mutants in WDR60 KO cells is shown in Fig. 7A. Both WT HA-WDR60 and HA-WDR60[T749M] efficiently rescued defects in the localization of IFT88 (Fig. 7B). However, expression of HA-WDR60[Q631*] was unable to restore the basal body localization of IFT88 (Fig. 7B and 7Bi). Similar data were obtained for the localization of IFT140 (Fig. 7C); expression of both WT and the HA-WDR60[T749M] mutant of WDR60 restored the localization of IFT140 to the base of the cilia, but in cells expressing HA-WDR60[Q631*], IFT140 was found throughout the cilium, as it was in WDR60 KO cells. Next, we tried to mimic the compound heterozygosity occurring in patient cells, generating a stable cell line that expresses both WDR60 [T749M] and [Q631*] mutants. When the two mutants were co-expressed in the same WDR60 KO cells we saw no additive effects or dominant negative effects, but cilia appeared normal with IFT88 only localized to the base (Fig. S3B, S3C) as was seen with the HA-WDR60[T749M] mutant.

**Figure 7:**
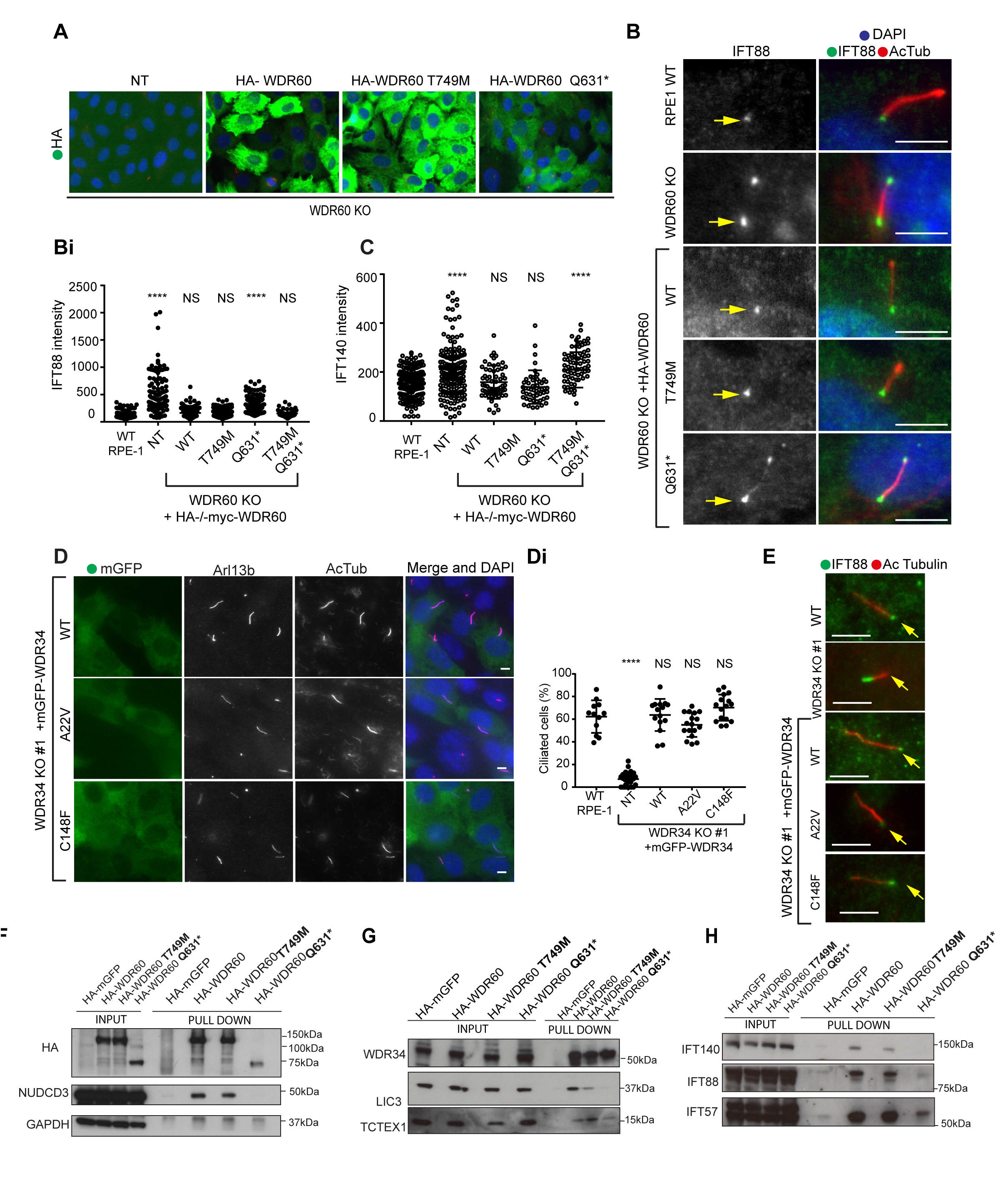
WDR34 and WDR60 KO rescue experiments. (A) Immunofluorescence staining of HA (green) in WDR60 KO cell lines stably expressing HA-tagged WT and mutant WDR60 constructs. (B and C) IFT88 or IFT140 staining (green) with AcTub labeling (red) of the stable cell lines shown in A as well as WT cells. (Bi) Total intensity quantification of IFT88 labeling across the length of primary cilia in each cell line (n=3; 97 WT, 125 WDR60 KO, 202 WDR60 KO+ HA-WDR60, 119 WDR60 KO+ HA-WDR60 [T749M], 150 WDR60 KO+ HA-WDR60 [Q631*] cells quantified). (Ci) Total intensity quantification of IFT140 labeling across the length of primary cilia in each cell line (n=3; 168 WT, 164 WDR60 KO, 63 WDR60 KO+ HA-WDR60, 56 WDR60 KO+ HA-WDR60 [T749M], 71 WDR60 KO+ HA-WDR60 [Q631*] cells quantified). (D and E) WDR34 KO#1 cells stably expressing mGFP-tagged WT and mutant WDR34. (D) Primary cilia staining with Arl13b (red) and AcTub (blue). (Di) Percentage of ciliated cells (n=3; 357 WT, 430 WDR34 KO, 399 WDR34 KO + mGFP-WDR34, 383 WDR34 KO + mGFP-WDR34 [A22V], 432 WDR34 KO + mGFP-WDR34[C148F] cells quantified). (E) IFT88 staining in WT, WDR34 KO#1 cells, and WDR34 KO#1 cells expressing GFP-tagged WT and mutant WDR34. One-way ANOVA followed by Kruskal-Wallis test was used p-value: ****=<0.0001. Arrows point to the ciliary base. (G-H) Immunoprecipitation of HA-tagged GFP, WT, WDR60 [T749M] and HA-WDR60 [Q631*] mutant proteins followed by immunoblot for (F) HA, NudCD3, and GAPDH; (G) WDR34, LIC3, and TCTEX1and (H) IFT140, IFT88, and IFT57.

In parallel, we also analyzed the phenotype of WDR34 KO cells stably expressing WT and mutant forms of WDR34. We found that expression of WT mGFP-WDR34 restored ciliogenesis and axoneme extension in WDR34 KO cells and that, unexpectedly, this was also true of cells expressing WDR34 incorporating either [A22V] or [C148F] mutations (Fig. 7D and 7Di). These disease-causing mutations were chosen as they lie in regions without specific domain prediction. The cilia that formed in WDR34 KO cells expressing either mGFP-WDR34[A22V] or mGFP-WDR34[C148F] were also positive for Arl13b (Fig. 7D). Moreover, WT and both WDR34 mutants were able to rescue IFT88 localization to the basal body (Fig. 7E).

To better understand the function of WDR60 we analyzed how the complex is assembled by performing immunoprecipitation of WT and mutant WDR60 proteins. We found that immunoprecipitation of HA-WDR60 expressed in WDR60 KO cells effectively pulls down the chaperone NudCD3, known to interact with dynein-2 via its WD repeat domains. As expected, NudCD3 did not bind to HA-WDR60[Q631*] lacking the WD repeat domain. WT WDR60 also bound effectively to WDR34 and, notably, this interaction was very similar with the HA-WDR60[T749M] or [Q631*] mutants (Fig. 7F). In contrast, LIC3 is readily detected with WT WDR60 but less so with WDR60[T749M]; only a very small amount of LIC3 was detected bound to WDR60[Q631*] (Fig. 7G). TCTEX1 was found to bind effectively to both WT and HA-WDR60[T749M] but less well to HA-WDR60[Q631*] (Fig. 8B). Next, we tested the interactions with IFT proteins, the primary cargo of the ciliary motors. We found that WT WDR60 can bind to IFT140, IFT88, and IFT57; WDR60[T749M] binds to all 3 IFT subunits tested but binds less well to IFT140 (Fig. 7H). In contrast, WDR60[Q631*] pulled down reduced levels of IFT88 and IFT57 and did not interact with IFT140.

**Figure 8:**
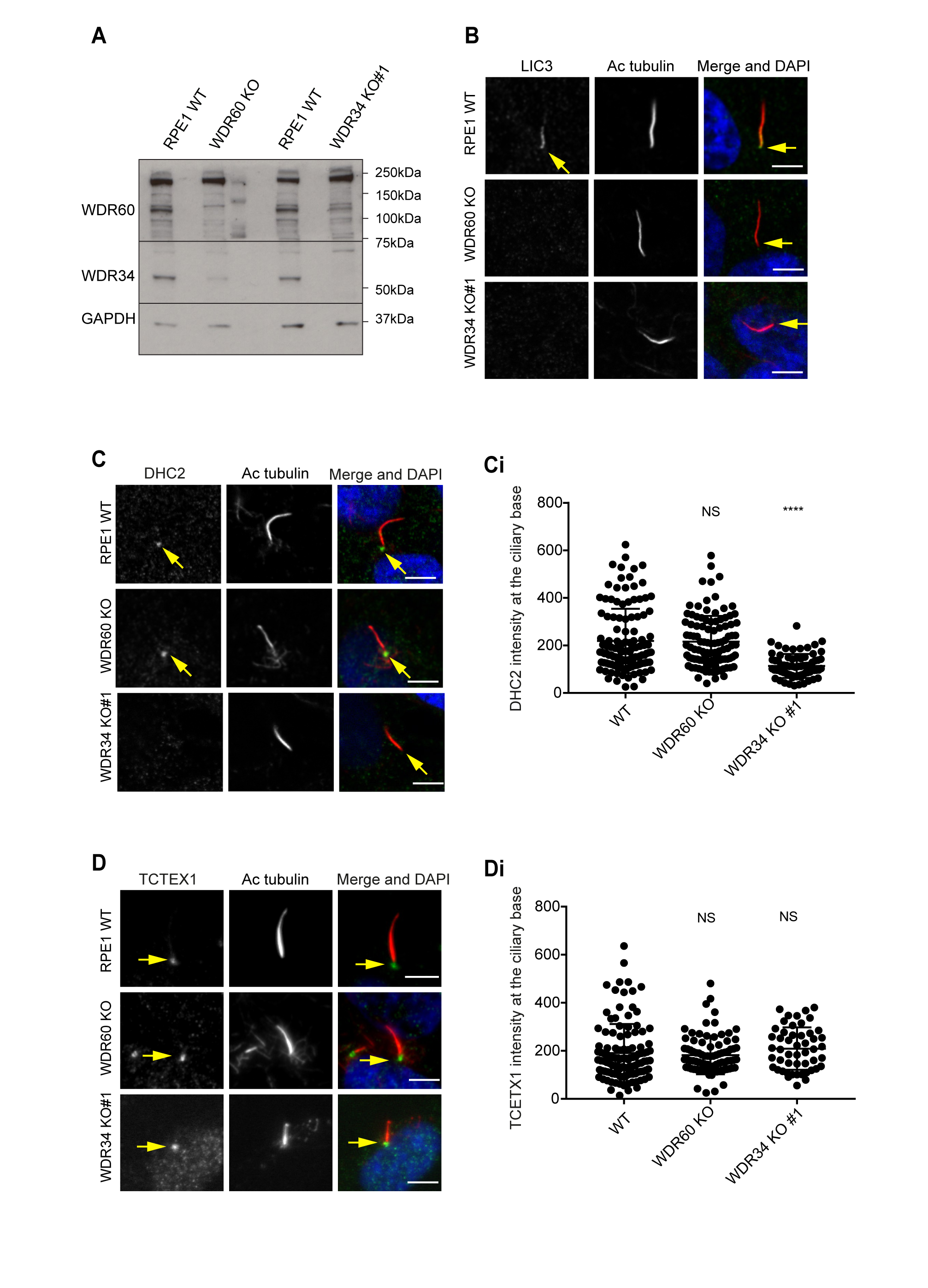
Dynein-2 assembly in primary cilium. (A) Immunoblotting for WDR60 and WDR34 in WT, WDR34 KO#1 and WDR60 KO cells. (B) LIC3 localization in the cilia of WT, WDR34 KO#1 and WDR60 KO cells. (C) DHC2 localization at the ciliary base in WT and KO cells. (Ci) Intensity quantification shows a reduction of DHC2 at the ciliary base in WDR34 KO#1 cells (n=3, 120 WT, 106 WDR60 KO and 71 WDR34 KO #1 cells quantified). (D) TCTEX1 localizes at the ciliary base in WT and KO cells (n=3 115 WT, 85 WDR60 KO, and 50 WDR34 KO#1 cells quantified). Mann-Whitney test, p-value: ****=<0.0001. Scale bars 5µm. Arrows point to the ciliary base.

### The stability of dynein-2 complex in WDR34 and WDR60 KO cells

It has been reported that loss of some components of dynein-2 modifies the stability of the whole dynein-2 complex. In *Chlamydomonas* depletion or loss of LIC3 (D1bIC2) causes a reduction of DHC2 in whole cells lysate, whereas the expression level of the intermediate chain is less affected (Reck et al., 2016). Similar results were obtained analyzing expression levels of DHC2 in patients cells with LIC3 mutations (Taylor et al., 2015). We have shown previously that siRNA depletion of WDR34 affects the stability of WDR60 and vice versa (Asante et al., 2014). To determine whether loss of one intermediate chain had an effect on the stability of the other, we analyzed levels of WDR34 and WDR60 in whole cells lysate of KO cells. Notably, we found that expression levels of WDR34 were reduced in WDR60 KO cells, although not completely lost. Correspondingly, there was a reduction of WDR60 expression levels in WDR34 KO whole cell lysate (Fig. 8A). Next, we sought to determine the effect of WDR60 and WDR34 loss on the localization of other dynein-2 subunits. We found that LIC3 (DYNC2LI1) localized in cilia of WT cells, but this localization was lost in WDR34 or WDR60 cells (Fig. 8B). DHC2 was detected at the base of the cilia in WT and WDR60 KO cells, but not along the cilia axoneme. Interestingly, DHC2 localization at the ciliary base was reduced in WDR34 KO cells (Fig. 8C). Moreover, we found that TCTEX1 (DYNC1LT1, Fig. 8D) was enriched at the base of the cilium in WT cells, with no changes in WDR34 or WDR60 KO cells. To test how the loss of one dynein-2 intermediate chain affected the localization of the other we overexpressed HA-WDR34 and HA-WDR60 in WDR60 and WDR34 KO cells. Both HA-WDR34 and HA-WDR60 were enriched at the base and in the cilia axoneme in WT cells (Fig. S4A and Fig. S4B). We observed no changes in the localization of HA-WDR34 in WDR60 KO cells, which was detected at the base and in the cilia (Fig. S4B). Surprisingly, HA-WDR60 was greatly enriched in stumpy cilia of WDR34 KO cells compared to the cilia in WT cells (Fig. S4A). Overexpression of HA-WDR60 could not rescue axoneme elongation in WDR34 KO cells (Fig. S4C). Additionally, overexpression of HA-WDR34 could not rescue abnormal IFT88 localization in WDR60 KO cells (Fig. S4D and S4Di).

### Defects in dynein-2 holocomplex assembly in absence of WDR34 or WDR60

The results described above suggest a defect in the assembly of the dynein-2 holoenzyme and therefore we used a proteomic approach to define the assembly of dynein-2. We stably expressed HA-WDR34 in WT and WDR60 KO cells and HA-WDR60 in WT and WDR34 KO cells and performed immunoprecipitations using HA-GFP as a control. Confirming that both WDR34 and WDR60 exist in the same complex, immunoprecipitation analysis showed that in WT cells HA-WDR60 pulls down WDR34, while HA-WDR34 pulls down WDR60 (Fig. S5A and S5B). Multiplex tandem-mass-tag (TMT) labeling enabled us to define the interactome of WDR34 in the presence and absence of WDR60 and of WDR60 in the presence and absence of WDR34. We found that the interactions of HA-WDR34 with dynein-2 components is reduced in WDR60 KO cells compared to WT cells (Fig. 9A). In particular, loss of WDR60 caused a reduction of DHC2, DYNLRB1, TCTEX1D2, DYNLT1, DYNC2LI1 and DYNLL2 immunoprecipitated with HA-WDR34 (Fig. 9B and 9Bi) compared to WT cells. Moreover, interactions of HA-WDR34 with several IFT components and the molecular chaperone NudCD3 were reduced in WDR60 KO cells. Surprisingly, HA-WDR60 interactions with dynein-2 components were largely unchanged between WT and WDR34 KO cells (Fig. 9B and 9Bii); dynein-2 components bound with similar efficiency to HA-WDR60 expressed either in WT and WDR34 KO cells, with a slight but reproducible reduction in DHC2 and DYNLRB1 binding in WDR34 KO compared to WT cells.

**Figure 9:**
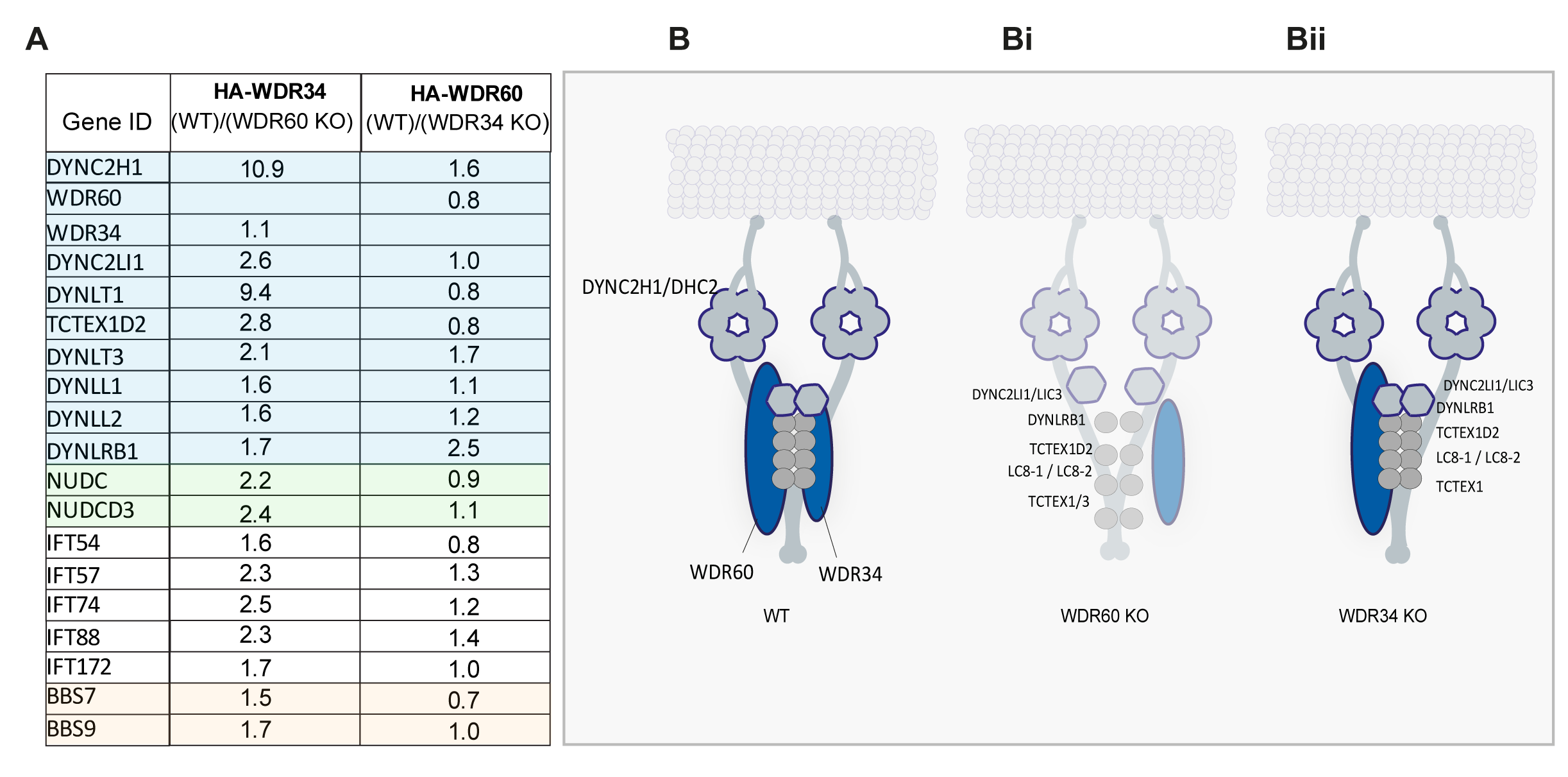
Proteomics of HA-WDR34 and HA-WDR60 interactomes in knockout cell lines. Abundance ratios in WT/ WDR34 KO cells expressing HA-WDR60 and WT/ WDR60 KO cells expressing HA-WDR34 determined by TMT analysis. Proteomic data were obtained from two independent experiments. The table shows raw data from one experiment. Similar results were obtained by normalizing the data with respect to the overexpressed protein abundance. (B) Schematic of the dynein-2 complex; the heavy chain is shown as two light blue homodimers, light intermediate chains are shown as hexagons, intermediate chains as extended ovals and light chains as circles. (Bi and Bii) Model of the dynein-2 complex in absence of (Bi) WDR60 or (Bii) WDR34. Reduced binding is represented by decreased color intensity. (Bi) In WDR60 KO cells interaction of HA-WDR34 with DHC2, DYNC2LI1/LIC3, DYNLT1/TCTEX1, DYNLL1/LC8-1, DYNLL2/LC8-2, TCTEX1D2, and DYNLRB1 is greatly reduced respect to WT cells. (Bii) In WDR34 KO cells interaction of HA-WDR60 with other components of dynein-2 is mostly unchanged.

## Discussion

### Structural and functional asymmetry of the dynein-2 motor

Our data provide evidence that the structural asymmetry with the dynein-2 motor is matched by functional asymmetry. Perhaps most strikingly, we find that WDR34 is essential for axoneme extension during early steps of ciliogenesis, whereas WDR60 is not required for ciliogenesis. Both subunits are necessary for maintaining proper cilia protein composition. Depletion of WDR34 using RNAi is also associated with ciliary defects (Asante et al., 2013) and patient fibroblasts have shorter cilia with a bulbous tip (Huber et al., 2013). Fibroblasts from WDR34 knockout mice also have stumpy cilia and defects in Shh signaling (Wu et al., 2017). It is intriguing that some cells missing WDR34 can still extend a rudimentary cilium but even here, ciliary protein localization is severely disrupted. Since WDR60 cells can extend an axoneme, WDR34 and WDR60 clearly have distinct but overlapping functions in cells. We consider that WDR34 plays an essential role in ciliogenesis to ensure delivery of a key factor required for axoneme extension. It is noteworthy that in the absence of WDR34, WDR60 can still assemble effectively with the other subunits of dynein-2. While these interactions are likely reduced compared to the normal situation, this shows that it is specifically WDR34 that is required at this early stage of ciliogenesis. Our EM data show that it acts at a stage after docking of the ciliary vesicle, immediately before axoneme extension. Paradoxically, our data also show that in the absence of WDR60, the dynein-2 holocomplex cannot form effectively yet axoneme extension occurs normally. This raises the possibility that WDR34 is itself required for axoneme extension, possibly outside of the context of the dynein-2 complex. We cannot rule out that there are dynein-2-independent functions of WDR34 and WDR60 but all data provide strong evidence that they co-exist in the dynein-2 holoenzyme.

### Assembly of the dynein-2 holocomplex

In WDR34 and WDR60 KO cells, LIC3 is no longer detected in primary cilia, while TCTEX1 localization, a dynein light chain that is also a component of dynein-1, was unperturbed. Coupled with our proteomics data, this suggests that the localization of LIC3 to cilia is a good reporter of dynein-2 assembly. Surprisingly we found that DHC2 levels at the ciliary base are reduced in WDR34 KO cells compared to WT and WDR60 KO cells. Our data show that in WDR34 KO cells, the remaining subunits can coassemble into partial dynein-2 complexes. We do not, however, know if this is a functional or indeed processive motor. According to the current model based on work in *C. elegans* and *Chlamydomonas*, the dynein-2 complex is transported by the kinesin-2 motor in the anterograde direction, as a passive cargo, and at the tip it switches direction and it is converted in an active motor (Chien et al., 2017; Mijalkovic et al., 2017; Pedersen et al., 2006; Toropova et al., 2017; Williamson et al., 2012). Within this model, it is not known how dynein-2 is maintained in an inactive state during anterograde trafficking. One possibility is that WDR34 might function in the final step of dynein-2 complex assembly prior to motor activation, potentially at the ciliary tip. Support for this comes from *C. elegans* where individual dynein-2 proteins show distinct turnaround times within cilia (Li et al., 2015). It is of course not clear how much of the protein is incorporated within a dynein complex in these experiments. The tight association of WDR34 with WDR60 and their functional interdependence (loss of one leading to destabilization of the other) would argue against the likelihood of dynamic subunit interchange during IFT. Further structural analysis could help resolve this question. Interaction of WDR60 with WDR34 in our pull-down experiments indicates that each dynein-2 complex contains both WDR34 and WDR60, in agreement with our own previously reported data (Asante et al., 2014). Moreover, we found that stably expressed WDR60 could not rescue ciliogenesis defects in WDR34 KO cells, neither could WDR34 rescue IFT88 localization defects observed in WDR60 KO cells. Therefore WDR34 and WDR60 are not functionally redundant, at least in this regard. This supports a model where WDR34 and WDR60 play different roles in ciliogenesis and in IFT within the context of the dynein-2 complex, likely through different interactions with distinct components.

### Ciliary trafficking defects in absence of WDR34 and WDR60

Loss of either WDR34 or WDR60 leads to IFT particle accumulation at the base of as well as within cilia. We found that loss of WDR60 results in an increase of IFT-B proteins, IFT20, IFT57, and IFT88, not only at the tip but also close to the cilia base. This suggests that IFT-B proteins could be retained at the basal body or around the transition zone. Consistent with these results, mutations in IFT-A or dynein-2 in mice also result in abnormal accumulation of IFT particles near the base of the cilium (Goggolidou et al., 2014; Liem et al., 2012; Ocbina et al., 2011). This has been linked to defects in the export of ciliary cargo across the transition zone (He et al., 2017). Similar defects are seen following disruption of the heavy chain, DHC2 (Hou and Witman, 2015).

In addition to defects in retrograde IFT, incomplete assembly of dynein-2 in WDR60 KO cells could cause a block in IFT train assembly and impaired entry of cargo into cilia. Indeed, work in C. *elegans* has shown that IFT-B is required for entry of dynein-2 into cilia (Yi et al., 2017). In addition to these defects, we show that KAP3, a subunit of kinesin-2, accumulates at the ciliary tip of WDR60 KO cells. This would decrease the levels of kinesin-2 available to load onto departing anterograde trains and further cause accumulation of IFT particles at the base. This might also reflect some functional coupling of dynein-2 and kinesin-2 in the assembly of anterograde IFT particles. This phenotype might be exacerbated because those IFT-B particles that do enter become stuck at the ciliary tips in WDR60 KO cells because of defects in retrograde IFT. Notably, our data are also consistent with models where, in metazoa, kinesin-2 motors are returned to the ciliary base by dynein-2-dependent retrograde IFT (Broekhuis et al., 2013; Chien et al., 2017; Mijalkovic et al., 2017; Prevo et al., 2015; Signor et al., 1999; Williams et al., 2014) but contrasts with *Chlamydomonas* where kinesin-2 diffuses back to the cilia base (Engel et al., 2012). Overall, our data support models where dynein-2 acts both in loading of cargo into cilia and in exit from cilia.

These findings suggest a complex interplay between IFT particles and IFT motors to control entry to and exit from cilia. Notably, interfering with either dynein-2 or IFT results in perturbation of transition zone organization, IFT, and ciliary membrane protein localization (Garcia-Gonzalo and Reiter, 2017).

### Dynein-2 is required for transition zone composition

The transition zone is the ciliary region most proximal to the mother centriole and it functions as a gate, acting as a diffusion barrier to prevent unregulated entry of high molecular weight proteins and maintaining the protein composition of the ciliary membrane (Garcia-Gonzalo and Reiter, 2017; Jensen and Leroux, 2017). Our results show that loss of either dynein-2 intermediate chain disrupts the organization of the ciliary transition zone. The defect is clearly very significant since we observe accumulation of intraciliary vesicles within the cilia of WDR60 KO cells that would normally be excluded by the diffusion barrier. The presence of intraciliary vesicles could also indicate the formation of ectosomes to remove excess membrane from the cilia. Such vesicles have been described in motile cilia (Shah et al., 2008) and in mouse photoreceptor cells (Gilliam et al., 2012) from BBS mutants, as well as in wild-type zebrafish (Goetz et al., 2014). However, we always see accumulated membrane at the tip of cilia in WDR60 KO cells when imaging living cells expressing ciliary membrane markers and have not detected any shedding of vesicles during such experiments. This suggests that any mechanisms to reduce membrane accumulation might not be able to overcome any defect in retrograde IFT. The presence of intraciliary vesicles in WDR60 KO cells is entirely consitent with a significant defect in the diffusion barrier and/or gated entry mechanisms. The role of dynein-2 in maintaining transition zone composition is also consistent with the fact that LIC3 and overexpressed WDR34 and WDR60 localize at the transition zone in RPE1 cells. Moreover, in a previous study, active phosphorylated TCTEX1 has been detected at the transition zone of neural progenitors (Li et al., 2011). Changes in transition zone composition are associated with a reduced localization of soluble and membrane proteins in the cilia (Berbari et al., 2008b; Chih et al., 2011). 5HT6, SSTR3, and Arl13B are all reduced in abundance within cilia of WDR60 KO cells, suggesting that these proteins might not be effectively retained within cilia, but leak out through the diffusion barrier. An alternative, as discussed above, is that these proteins are less effectively loaded into cilia. We did not find a difference in the intensity levels of overexpressed Rab8a in WDR60 KO compared to WT cells suggesting that at least some proteins can enter normally. It seems likely that the defects we see in both entry to, and exit from, cilia in these KO cells are caused by defects in transition zone structure.

### Dynein-2 dysfunction and disease

The list of mutations causing disease in genes associated with primary cilia is continuously expanding. Our results show that Smo localization is deregulated in the absence of WDR60. Smo is thought to enter cilia continuously but rapidly exit in the absence of ligand. Our data are consistent with this, with loss of WDR60 results in aberrant accumulation of Smo in cilia and likely to perturbed hedgehog signaling. Interestingly, abnormal accumulation of Smo in the cilia has also been observed in mouse fibroblasts with mutations in DHC2. In these mutants mice inactivation of dynein-2 causes loss of Shh signaling and midgestation lethality (Ocbina et al., 2011). Given the links between hedgehog signaling and skeletogenesis, this is likely to be a major cause of the phenotypes seen on loss-of-function of dynein-2 in animal models and in patients (Dagoneau et al., 2009; Li et al., 2015; May et al., 2005; Wu et al., 2017).

In this study, we also characterized the function of dynein-2 using disease-causing mutations found in SRPs and JATD syndromes. Using mutagenesis to recreate patient mutations WDR60[Q631*] and WDR60[T749M], we show that the N-terminal region of WDR60 is sufficient to bind to WDR34 and TCTEX1 but not to LIC3. This suggests some assembly of a WDR34-WDR60 module prior to full complex assembly. Further analysis of patient mutation [Q631*] reveals that the C-terminal ß-propeller domain of WDR60 is required for binding of the IFT-B proteins, including IFT88. The observed reduction in binding between the WDR60[Q631*] mutant and NudCD3 is expected as the NudC family acts as co-chaperones with hsp90 for folding of β-propellers such as WD repeat domains (Taipale et al., 2014). In contrast, WDR60[T749M] binds to other dynein-2 proteins, IFT proteins, and NudCD3. The reduction in binding to LIC3 seen with this mutation suggests that less efficient dynein-2 assembly might contribute to the patient phenotypes.

Surprisingly, expression of WT and WDR34[A22V] or [C148F] mutants in WDR34 KO cells was able to rescue both normal cilia length and normal IFT88 localization at the cilia base. Genome sequencing data showed that patients with WDR34[A22V] mutation also present mutations in DYNC2H1 and IFT140 (Schmidts et al., 2013b), similarly, a second mutation in the WDR34 gene was found in patients with [C148F] mutant. Thus, it is possible that these two mutant proteins are not sufficient to impair dynein-2 function themselves and exclusively cause a deleterious effect when a second modification is also present.

Our data show that not only is dynein-2 required for retrograde IFT but also to build and maintain a functional diffusion barrier at the base of the cilium. Our data do not discriminate between roles in assembly versus maintenance of the transition zone. Other recent work has shown that Joubert syndrome is caused by disruption of the transition zone (Shi et al., 2017). Joubert syndrome leads to severe neurological effects that are underpinned by developmental defects in hedgehog signaling. Jeune syndrome is characterized by skeletal defects but these are attributed to hedgehog signaling defects. Together our work and that of Shi et al. (2017) suggests that defects in transition zone architecture and resulting defects in developmental signaling might define a common root cause of both Joubert and Jeune syndromes, and indeed perhaps other ciliopathies.

## Acknowledgements

We would like to thank Katharine Risk, Beth Moyse, and Imogen Binnian for their contributions to the work and Janine McCaughey for helpful discussion on this work. Thanks also to Max Nachury and Jackie Goetz for helpful comments. This work was supported by grants from the BBSRC (BB/N000420/1) and MRC (MR/P000177/1 and MR/K018019/1). We would like to thank the Wolfson Bioimaging Facility for support of the microscopy experiments. Confocal microscopy was supported by a BBSRC ALERT 13 capital grant (BB/L014181/1).

The authors declare no competing financial interests.

Author contributions: DJS, LV, and NLS conceived and designed the experiments and wrote the manuscript. LV and NJS performed the experiments and analyzed the data. KJH helped design and conducted the proteomics experiments and data analysis.

## Materials and Methods

All reagents were purchased from Sigma-Aldrich (Poole, UK) unless stated otherwise.

### Plasmids, cloning, and mutagenesis

The human WDR34 gene was obtained from the Origene (SC319901, Cambridge Bioscience), human WDR60 was generated by gene synthesis (Life Technologies, Paisley, UK). An HA tag for WDR34 and WDR60 was added by PCR and both proteins were subcloned in the pLVX-puro vector. Mutant [T749M WDR60] was generated by site-directed mutagenesis PCR using the primers: Fw: 5’-CAGAACCGCCatgTTCTCCACC-3’ and Rv 5’-GGTGGAGAACATGGCGGTTCTG-3’ changing codon ACG (Threonine) to ATG (Methionine). WDR60 [Q631*] mutant was constructed by site-directed mutagenesis using the primers: Fw: 5’-GATAGCAGCTCCtagCTGAATACC-3’ and 5’-GGTATTCAGCTAGGAGCTGCTATC-3’ changing codon CAG (Glutamine) to TAG (STOP codon). For WDR34, the A22V mutant was generated by changing codon GCG (Alanine) to GTC (Valine) using oligonucleotides 5’-TGCGGCGCTGgtgACAGTCGGGG-3’ and 5’-ACACCAGCGCTTCCCGCCTG-3’. The C148F mutation was generated by changing codon TGT (Cysteine) to TTT (Phenylalanine) using oligonucleotides 5’-GATGGTGTCTtttCTGTATACCCTGGG-3’ and 5’-TGCTGCTGCTCGGTCCAG-3’. All constructs were validated by DNA sequencing.

Mouse L13-Arl13b-GFP was a gift from Tamara Caspary (Addgene plasmid # 40879, (Larkins et al., 2011)), IFT20-GFP ((Follit et al., 2006), plasmid JAF2.13) was a gift from Gregory Pazour (Addgene plasmid # 45608). pEGFPN3-SSTR3 and pEGFPN3-5HT6 were gifts from Kirk Mykytyn (Addgene plasmid #35624 and #35623, (Berbari et al., 2008a)), EGFP-Rab8 was a gift from Johan Peränen (University of Helsinki).

### Cell culture

Human telomerase-immortalized retinal pigment epithelial cells (hTERT-RPE-1, ATCC CRL-4000) were grown in DMEM-F12 supplemented with 10% FBS (Life Technologies, Paisley, UK) at 37°C under 5% CO2. Cells were not validated further after purchase from ATCC. Transient transfections of Arl13b-GFP, SSTR3-GFP, 5HT6-GFP, and Rab8-GFP were performed using Lipofectamine 2000 (Life Technologies, Paisley, UK) according to the manufacturer's protocol. Lentiviral particles for each of the stable RPE-1 cell lines were produced in HEK293T cells using the Lenti-XTM HTX Packaging System (Clontech, Saint-Germain-en-Laye, France). Low passage hTERT-RPE1 cells were transduced with the resultant viral supernatant, strictly according to the manufacturer's directives and at 48 h post-transduction, cells were subcultured in presence of 5 µ.g/ml puromycin. RPE-1 cells were incubated in serum-free medium for 24 h to induce ciliogenesis. Confluent cells were placed in serum-free media and treated with Shh agonist SAG (Selleckchem (from Stratech Scientific, Ely, UK) Catalog No.S7779) at the final concentration of 100 nM for 24 h.

### Genome engineering

The guide RNAs (gRNA) targeting WDR34 were designed using ‘chop chop’ software (Labun et al., 2016) or using CRISPR design http://crispr.mit.edu/ for designing WDR60 gRNA (Hsu et al., 2014). pSpCas9(BB)-2A-GFP (Addgene plasmid, #PX458) was used as the vector to generate a gRNA. The gRNA sequences (5’-A GCC TTT CTT CGG AGA GTG G-3’; and 5’-CA GGT GTC TTG TCT GTA TAC -3’) were designed to target Exon2 and Exon3 of human WDR34. Similarly, the gRNA (5’-AG GTG CAG GGA TCC CGA CCA-3’) was designed to target exon 3 of WDR60. RPE-1 cells were transfected with 1 u.g of pSpCas9(BB)-2A-GFP. After 48 h GFP-positive cells were sorted, and singles cells were plated in a 96 well plate. To check the WDR34 and WDR60 genes, genomic DNAs of the target sequences were extracted and subjected to PCR. Subsequently, the PCR products were cloned in the pGEM^®^ T Easy vector according to the manufacturer's instructions and sequenced. In three cells clones, identified as WDR34 KO#1, WDR34 KO#2 and WDR60 KO, small deletions/insertions causing a frameshift were detected in both alleles (Supplementary Fig. 1 for details). On the contrary, the cell clones identified as CRISPR CTRL WDR34 and WDR60 cells, transfected and treated in the same conditions of our knock out clones, did not show any mutation in the targeted genomic DNA region.

### Antibodies

The antibodies used, and their dilutions for western blotting (WB) and immunofluorescence (IF) are as follows: Acetylated tubulin (Sigma (Poole, UK) T6793 1:2000 for IF), rabbit anti-HA (Cell Signaling Technologies (New England Biolabs, Hitchin, UK) 1:2000 WB, 1:1000 IF), rabbit IFT88 (Proteintech (Manchester, UK) 13967-1-AP, 1:200 WB, 1:300 IF), rabbit anti-IFT140 (Proteintech 17460-1-AP, 1:200, WB 1:100 IF), rabbit anti-IFT57 (Proteintech 11083-1,-AP 1:200, WB 1:100 IF), rabbit anti-IFT43 (Proteintech 24338-1-AP, 1:50 IF), anti-IFT20 (Proteintech 13615-1-AP, 1:200 IF), anti-TMEM67 (proteintech13975-1-AP, 1:50 IF), anti-RPGRIP1L (Proteintech 55160-1-AP, 1:100 IF), anti-DYNC2HC1 (Proteintech 55473-1-AP, 1:100 IF), rabbit anti-LIC3 (Proteintech 15949-1-AP, 1:250 WB, 1:100 IF), rabbit anti-TCTEX1 (Santa Cruz Biotechnology (from Insight Biotechnology, Wembley, UK) sc-28537, 1:200 WB, 1:100 IF), rabbit anti-Arl13B (Proteintech 17711-1AP, 1:1000 IF), rabbit anti-TCTN1 (Proteintech 15004-1-AP, 1:100 IF), rabbit anti-Smo (Abcam (Cambridge, UK) ab38686, 1:100 IF), Sheep anti-Myc ((Fan et al., 2010) kindly provided by Harry Mellor, University of Bristol), rabbit anti-WDR60 (Novus Biologicals (from Bio-Techne Abingdon, UK) NBP1-90437 1:300 WB in Fig.9), rabbit anti-WDR60 (Sigma HPA021316, 1:300 WB in Fig. S2 and Fig.S5), rabbit anti-WDR34 (Novus NBP188805, 1:300 WB), mouse anti-GAPDH (Abcam ab9484, 1:1000 WB), p150 glued (BD 6127009, 1:1000 WB), LIS1 (Bethyl A300-409A, 1:1000 WB), dic74 (MAB1618 Millipore (Watford, UK), 1:1000 WB), NUDCD3 (Sigma HPA019136, 1:350 WB).

### Immunofluorescence

Cells grown on 0.17 mm thick (#1.5) coverslips (Fisher Scientific, Loughborough, UK) were washed in PBS, and then fixed for 10 min in PFA and permeabilized with PBS containing 0.1% Triton X-100 for 5 min. Alternatively, cells were fixed in ice-cold methanol at -20°C for 5 min for TMEM67, RPGRIP1L, TCTN1, DYNC2H1 and IFT20 staining.

For TCTEX1 immunolabelling, cells were washed twice with pre-warmed cytoskeletal buffer (CB, containing 100 mM NaCl 300 mM sucrose, 3 mM MgCl2, and 10 mM PIPES) and fixed for 10 min in CB-PFA, as described previously (Hua and Ferland, 2017).Subsequently, cells were blocked using 3% BSA in PBS for 30 min at room temperature. The coverslips were incubated with primary antibodies for 1 h at room temperature, washed in PBS and then incubated with secondary antibodies for another 1 h at room temperature. Nuclear staining was performed using DAPI [4,6-diamidino-2-phenylindole (Life Technologies), diluted at 1:5000 in PBS] for 3 min at room temperature, and the cells were then rinsed twice in PBS. Cells were imaged using an Olympus IX-71 or IX-81 widefield microscope with a 63x objective, and excitation and emission filter sets (Semrock, Rochester, NY) controlled by Volocity software (version 4.3, Perkin-Elmer, Seer Green, UK). Alternatively, cells in Fig. 4 and Fig. 7 were imaged using Leica SP5 confocal microscope (Leica Microsystems, Milton Keynes, UK). Live images in Fig. 3 were imaged using Leica SP8. All images were acquired as 0.5 µ.m z-stacks. All graphs show mean and standard deviation.

### Rescue experiments

For ‘rescue’ experiments, stable WDR60 KO cell lines overexpressing wild-type and mutants HA-tagged WDR60 were generated. Similarly, WDR34 KO#1 cells were stably transfected with WT and mutants WDR34 tagged with a GFP. Cells were serum starved for 24 h, fixed and processed for immunofluorescence analysis.

### Electron microscopy

Cells were serum starved 24 h and fixed in 2.5% glutaraldehyde for 20 min. Next, the cells were washed for 5 min in 0.1 M cacodylate buffer then post-fixed in 1% OsO4/0.1 M cacodylate buffer for 30 min. Cells were washed 3x with water and stained with 3% uranyl acetate for 20 min. After another rinse with water, cells were dehydrated by sequential 10 min incubations with 70, 80, 90, 96, 100 and 100% ethanol before embedding in Epon™ at 70°C for 48 h. Thin (70 nm) serial sections were cut and stained with 3% uranyl acetate then lead citrate, washing 3x with water after each. Once dried, sections were imaged using an FEI (Cambridge, UK) Tecnai12 transmission electron microscope.

### Immunoblotting

Cells were lysed in buffer containing 50 mM Tris pH7.5, 150 mM NaCl, 1% Igepal and 1 mM EDTA. Samples were separated by SDS-PAGE followed by transfer to nitrocellulose membranes. Membranes were blocked in 5% milk-TBST. Primary antibodies diluted in blocking buffer were incubated with membrane overnight and detected using HRP-conjugated secondary antibodies (Jackson ImmunoResearch, West Grove, PA) and enhanced chemiluminescence (GE Healthcare, Cardiff, United Kingdom).

### Fluorescence intensity measurement

Quantification of fluorescence intensity was performed using original images. Measurement of intensity was performed using the average projections of acquired z-stacks of the area of the ciliary marker acetylated tubulin. Fluorescence intensity along the ciliary axoneme was measured using ImageJ plot profile tool. Fluorescence intensity in at the ciliary base was measured drawing same diameter circles at the ciliary base.

### Immunoprecipitation

RPE-1 cells expressing the indicated cDNA constructs were washed with PBS and incubated with the crosslinker solution (1 mM DSP, Thermo Fisher Scientific #22585) for 30 min on ice. The reaction was quenched by adding 500 mM Tris-HCl pH 7.5 for 15 min. Cells were washed twice with PBS and lysed in a buffer containing 50 mM Tris/HCl, pH 7.4, 1 mM EDTA, 150 mM NaCl, 1% Igepal and protease inhibitors (539137, Millipore). Subsequently, cells were incubated on a rotor at 4°C for 30 min and then lysates were centrifuged at 13,000 g at 4°C for 10 min. Cell lysates were added to the equilibrated anti-HA-Agarose beads (Sigma A2095, batch number 026M4810V) and incubated on a rotor at 4°C. Next, the beads were washed three times by centrifuging at 2000 g for 2 min at 4°C with 1 ml of washing buffer (50 mM Tris-HCl, 150 mM NaCl, 0.5 mM EDTA, Triton X-100 0.3% SDS 0.1%) containing protease inhibitors (539137, Millipore). Samples used for SDS-PAGE and immunoblotting were resuspended in 50 µ.l of LDS sample buffer (Life Technologies) containing sample reducing agent (Life Technologies) and boiled at 95°C for 10 min.

### Proteomic analysis

For TMT Labelling and high pH reversed-phase chromatography, the samples were digested from the beads with trypsin (2.5 µ.g trypsin, 37°C overnight), labeled with Tandem Mass Tag (TMT) six-plex reagents according to the manufacturer's protocol (Thermo Fisher Scientific, Loughborough, UK) and the labeled samples pooled. The pooled sample was then desalted using a SepPak cartridge according to the manufacturer's instructions (Waters, Milford, Massachusetts, USA)). Eluate from the SepPak cartridge was evaporated to dryness and resuspended in buffer A (20 mM ammonium hydroxide, pH 10) prior to fractionation by high pH reversed-phase chromatography using an Ultimate 3000 liquid chromatography system (Thermo Fisher Scientific). In brief, the sample was loaded onto an XBridge BEH C18 Column (130 Å, 3.5 µm, 2.1 mm X 150 mm, Waters, UK) in buffer A and peptides eluted with an increasing gradient of buffer B (20 mM ammonium hydroxide in acetonitrile, pH 10) from 0-95% over 60 min. The resulting fractions were evaporated to dryness and resuspended in 1% formic acid prior to analysis by nano-LC MSMS using an Orbitrap Fusion Tribrid mass spectrometer (Thermo Fisher Scientific).

### Nano-LC Mass Spectrometry

High pH RP fractions were further fractionated using an Ultimate 3000 nano-LC system in line with an Orbitrap Fusion Tribrid mass spectrometer (Thermo Fisher Scientific). In brief, peptides in 1% (vol/vol) formic acid were injected onto an Acclaim PepMap C18 nano-trap column (Thermo Fisher Scientific). After washing with 0.5% (vol/vol) acetonitrile 0.1% (vol/vol) formic acid, peptides were resolved on a 250 mm x 75 µm Acclaim PepMap C18 reverse phase analytical column (Thermo Fisher Scientific) over a 150 min organic gradient, using 7 gradient segments (1-6% solvent B over 1 min., 6-15% B over 58 min., 15-32% B over 58 min., 32-40% B over 5 min., 40-90% B over 1 min., held at 90% B for 6 min and then reduced to 1% B over 1 min.) with a flow rate of 300 nl min-1. Solvent A was 0.1% formic acid and Solvent B was aqueous 80% acetonitrile in 0.1% formic acid. Peptides were ionized by nano-electrospray ionization at 2.0 kV using a stainless steel emitter with an internal diameter of 30 µm (Thermo Fisher Scientific) and a capillary temperature of 275°C.

All spectra were acquired using an Orbitrap Fusion Tribrid mass spectrometer controlled by Xcalibur 2.0 software (Thermo Fisher Scientific) and operated in data-dependent acquisition mode using an SPS-MS3 workflow. FTMS1 spectra were collected at a resolution of 120 000, with an automatic gain control (AGC) target of 400 000 and a max injection time of 100 ms. Precursors were filtered with an intensity range from 5000 to 1E20, according to charge state (to include charge states 2-6) and with monoisotopic precursor selection. Previously interrogated precursors were excluded using a dynamic window (60 s +/- 10 ppm). The MS2 precursors were isolated with a quadrupole mass filter set to a width of 1.2 m/z. ITMS2 spectra were collected with an AGC target of 10 000, max injection time of 70 ms and CID collision energy of 35%.

For FTMS3 analysis, the Orbitrap was operated at 30 000 resolution with an AGC target of 50 000 and a max injection time of 105 ms. Precursors were fragmented by high energy collision dissociation (HCD) at a normalized collision energy of 55% to ensure maximal TMT reporter ion yield. Synchronous Precursor Selection (SPS) was enabled to include up to 5 MS2 fragment ions in the FTMS3 scan.

### Data Analysis

The raw data files were processed and quantified using Proteome Discoverer software v2.1 (Thermo Fisher Scientific) and searched against the UniProt Human database (140000 entries) and GFP sequence using the SEQUEST algorithm. Peptide precursor mass tolerance was set at 10 ppm, and MS/MS tolerance was set at 0.6 Da. Search criteria included oxidation of methionine (+15.9949) as a variable modification and carbamidomethylation of cysteine (+57.0214) and the addition of the TMT mass tag (+229.163) to peptide N-termini and lysine as fixed modifications. Searches were performed with full tryptic digestion and a maximum of 1 missed cleavage was allowed. The reverse database search option was enabled and the data was filtered to satisfy false discovery rate (FDR) of 5%.

